# Tomato drought-responsive transcription factor TINY1 suppresses embryonic growth

**DOI:** 10.1101/2025.10.22.683859

**Authors:** Matar Azriel, Hagai Shohat, Dalia Blinderman, David Weiss, Yotam Zait

**Affiliations:** Institute of Plant Sciences and Genetics in Agriculture, The Hebrew University of Jerusalem, P.O. Box 12, Rehovot 76100, Israel

**Author notes:** Correspondence: Yotam Zait, David Weiss Email addresses. Matar Azriel -, Hagai Shohat-, Dalia Blinderman. Highlight- The tomato drought-induced DREB transcription factor TINY1 suppressed embryonic and early seedling growth by inhibiting gibberellin activity, but it had no significant effect on drought response mechanisms.

**Keywords:** DREB, drought, embryo growth, gibberellin, TINY1, Tomato, transpiration

## Abstract

Dehydration-Responsive Element-Binding (DREB) transcription factors play an important role in plant responses to drought. DREB subfamily A4, contains a sub-group named TINY. Previous studies in Arabidopsis suggest that TINYs suppress plant growth and mediate abscisic acid (ABA)-induced stomatal closure. In this study, we investigated the function of the tomato drought-induced *TINY1*. Under drought conditions, *tiny1* mutant lost turgor and wilted more rapidly than control M82 plants. However, this sensitivity was attributed to its larger leaf area, rather than intrinsic differences in drought response. Measurements of stomatal conductance, leaf temperature, and osmotic adjustment revealed no significant differences between *tiny1* and M82. Furthermore, whole plant daily transpiration of M82 and *tiny1* with similar leaf area, showed no differences. Interestingly, the growth-promoting effect of *tiny1* was confined to early developmental stages; enhanced embryo growth and hypocotyl elongation, and accelerated emergence of the first true leaves—trait that later contributed to increased leaf area. At later stages, the mutation had no observable impact on growth rate. Our results show increased gibberellin (GA) activity in the mature *tiny1* embryo and suggest that TINY1 suppresses embryonic growth by repressing GA biosynthesis through downregulation of *GA 20-oxidase 4* (*GA20ox4*) gene expression.

## Introduction

Water deficiency suppresses directly and indirectly major biochemical pathways in plants, including photosynthesis and primary carbon metabolism (Tardieu *et al*., 2018). It also inhibits plant growth, flowering and fruit development (Gupta *et al*., 2020). Plants use two major strategies to cope with drought: drought avoidance and drought tolerance (Skirycz and Inze, 2010; Kooyers, 2015). ’Drought avoidance’ is a major plant adaptation strategy to survive transient water-deficit conditions. To avoid drought stress, plants reduce their transpiration and can use the available water in the soil more slowly and for a longer period before the arrival of the next rain. Several mechanisms have evolved to reduce water loss under drought, including fast stomatal closure and long-term growth inhibition (Kalladan *et al*., 2017). Stomatal closure is regulated mainly by abscisic acid (ABA), which accumulates in response to drought and triggers a signaling cascade that leads to ion efflux, loss of guard cell turgor, and stomatal pore closure (Bharath *et al*., 2021). Growth inhibition is regulated by the reduced turgor pressure, which is the driving force for cell expansion (Ray *et al*., 1972;; Ali, 2023), reduced levels of growth promoting hormones, such as gibberellin (GA) and increased activity of ABA (Shohat *et al*., 2021). Tolerance to drought is acquired mainly through osmotic adjustment (Blum, 2017), scavenging of reactive oxygen species (ROS) (Noctor *et al*., 2014), activation of stress-related genes and accumulation of drought related proteins (Zhang *et al*., 2022).

The Dehydration-Responsive Element-Binding (DREB) family of transcription factors play a crucial role in plant responses to various abiotic and biotic stresses, including drought. These proteins belong to the APETALA2/Ethylene Responsive Factor (AP2/ERF) superfamily and are characterized by their ability to bind to specific DNA sequences called DRE or CRT elements (Agarwal *et al*., 2006). DREBs affect and regulate various physiological and developmental stress responses, including stomatal closure, growth suppression and induction of stress-related genes (Lata and Prasad, 2011; Sarkar *et al*., 2019). DREB proteins are divided into six subfamilies (A1 to A6). Subfamily A4, contains 16 genes in Arabidopsis, including a sub group named TINY (Nakano *et al*., 2006). Arabidopsis TINY1, a member of this family, is a unique transcription factor that connects both abiotic and biotic stress signaling pathways. It binds to both DRE and Ethylene-Responsive Element (ERE) sequences, making it a versatile regulator of stress responses in plants (Sun *et al*., 2008). TINY was first identified in an activation-tagging screen for gain-of-function mutations in Arabidopsis (Wilson *et al*., 1996). The mutant is dwarf due to a reduction in cell expansion and therefore named TINY. TINY genes are induced by various abiotic stresses, including drought, cold and salt. The triple Arabidopsis mutant *tiny1tiny2tiny3* has larger leaves and longer petioles (Xie *et al*., 2019), whereas TINY1 overexpressing plants have smaller leaves, increased drought-responsive gene expression, and they are hypersensitive to ABA-mediated stomatal closure (Sun *et al*., 2008; Xie *et al*., 2019). The effect of TINY on growth and drought response in Arabidopsis was associated with its interaction with the brassinosteroid (BR) pathway (Xie *et al*., 2019).

Tomato has six putative DREB-TINY genes and drought conditions induced the expression of *TINY1* (Shohat *et al*., 2021). *SlDREB* (TINY1 in Shohat *et al*., 2021) overexpression suppresses the expression of the gibberellin (GA) biosynthesis genes *ent-copalyl diphosphate synthase* (*CPS*), *GA 20-oxidase1* (*GA20ox1*), *GA20ox2* and *GA20ox4*, reduced GA levels, inhibits growth and promotes drought resistance (Li *et al*., 2012). Our previous study in tomato, using *tiny1* CRISPR mutant shows that the loss of TINY1 activity has no effect on *GA20ox1* and *GA20ox2* expression, but attenuated the induction of the GA deactivating gene *GA2ox7* by drought (Shohat *et al*., 2021). Thus, the role of TINY1 in the regulation of GA accumulation under drought is not yet clear.

Here, we demonstrate that the loss of TINY1 activity in tomato promoted embryonic growth, hypocotyl elongation, and the formation of the first true leaf. However, beyond these early stages, *tiny1* did not influence growth rate under either well-watered or drought conditions. Our results suggest that the loss of TINY1 increases GA activity in the growing embryo, probably due to the elevated expression of *GA20ox4*. Under drought conditions, *tiny1* mutants lost turgor more rapidly than the wild type (WT), but this was not attributable to delayed stomatal closure or increased stomatal conductance but was due to the presence of an additional leaf which was produced early in seedling development but resulted later in a larger total leaf area.

## Materials and Methods

### Plant materials and growth conditions

The loss-of-function CRISPR-Cas9 derived *tiny1* alleles #1 (Shohat *el al.,* 2021) and #23 and the transgenic line *35S:proΔ17* (Nir *et al*,.2017) are all in the M82 background. The *tiny1*#23 allele is a newly identified line not reported previously. This allele has an 89-base pair deletion and an early stop codon after 270 bp. (Supplementary Fig. S1). Plants were grown in a growth room set to a photoperiod of 12 h:12 h, day:night, light intensity of 150 μmol m^−2^ s^−1^, and room temperature of 25°C. In other experiments, plants were grown in a greenhouse under natural day-length conditions, a light intensity of 700 - 1000 µmol m^−2^ s^−1^ and a temperature of 18–30°C. Seeds were harvested from ripe fruits taken from plants grown in greenhouse, thoroughly washed with water, and incubated overnight at 37°C in 10% sucrose solution, then rinsed and surface-sterilized using 1% sodium hypochlorite followed by a wash with 1% Na_3_PO_4_·12H_2_O. Sterilized seeds were dried and stored at room temperature until use.

### Drought treatments

Plants were initially irrigated to saturation, after which irrigation was withheld to induce drought stress. Leaf relative water content (RWC; see below) was assessed.

### Measurements of RWC

Leaf RWC of irrigated and drought-treated plants were measured as follows: Fresh weight (FW) was measured immediately after leaf detachment and then leaves were soaked for 24 h in 5 mM CaCl2 and the turgid weight (TW) was recorded. The leaves were subsequently dried at 55°C for 72 hours to obtain the dry weight (DW). Leaf RWC was calculated using the following formula:

Leaf RWC (%) = [(FW − DW) / (TW − DW)] × 100.

### Stomatal conductance (*g*_s_) measurements

Stomatal conductance (gs) was measured on the terminal leaflet of the third fully expanded leaf (from the shoot apex) using an LI-600 Porometer (LI-COR Biosciences, Lincoln, NE, USA). All measurements were taken between 09:00 and 11:00 under controlled growth room conditions on 4-week-old plants.

### Thermal imaging

Thermal images were obtained using an A655sc, infrared camera with a 15° field of view (FLIR Systems, Wilsonville, OR, USA). The camera was mounted vertically above the plants to capture the entire canopy. Mean leaf temperature was calculated for the total leaf area using a customized region of interest (ROI) tool, following the manufacturer’s instructions.

### Leaf sap osmolality measurements

Terminal leaflets from the third fully expanded leaf (counting from the shoot apex) were collected, immediately frozen in liquid nitrogen and stored at −80°C. For sap extraction, samples were rapidly thawed at 37°C and transferred into 1.5 mL Eppendorf tubes with perforated bottoms. These tubes were placed inside larger collection tubes and centrifuged at 12,000 rpm for 10 minutes to collect the sap. A 25 µL aliquot of the extracted sap was used to measure osmolyte concentration using a freezing-point depression (http://www.loeser-osmometer.de/home-eng.html) osmometer (Basic M, Löser Messtechnik, Germany).

### Whole-plant daily transpiration measurement

Whole-plant transpiration rate was measured using a high-throughput telemetric, gravimetric-based phenotyping system (Plantarray 3.0 system; Plant-DiTech, Israel), as described by Dalal *et al*. (2020). The experiment was conducted in the lysimeter greenhouse of the I-CORE Center for Functional Phenotyping at the Hebrew University (Rehovot, Israel) (http://departments.agri.huji.ac.il/plantscience/icore.phpon) under three irrigation regimes: control (full irrigation), moderate drought (irrigation at 50% of daily transpiration), and terminal drought (no irrigation). Plants were grown in 3.5-L pots under semi controlled environmental conditions with day/night temperatures of 30°C/18°C and exposed to the natural photoperiod and ambient light intensity (700 - 1000 µmol m^−2^ s^−1^).

Each pot was placed on a load cell in a randomized block arrangement and the soil surface was sealed to prevent evaporation. The data were analyzed using SPAC Analytics (Plant-Ditech, Yavne, Israel) software to obtain the following whole-plant physiological traits: daily transpiration, calculated as the weight loss between predawn and sunset and transpiration rate, calculated as the weight loss between two 3-min time points.

### Measurements of leaf area

Total leaf area per plant was measured using a LI-3100 leaf area meter (LI-COR Biosciences, Lincoln, NE, USA).

### RNA extraction and cDNA synthesis

Total RNA was extracted from frozen plant tissues-including red ripe fruit seeds, hypocotyls, seedlings, or terminal leaflets of the third fully expanded leaf-using the RNeasy Plant Mini Kit (Qiagen, Hilden, Germany). For cDNA synthesis, 3 µg of total RNA was reverse transcribed using SuperScript II reverse transcriptase (18064014; Invitrogen, Waltham, MA, USA), following the manufacturer’s instructions.

### RT-qPCR analysis

RT-qPCR analysis was performed using an Absolute Blue qPCR SYBR Green ROX Mix (AB-4162/B) kit (Thermo Fisher Scientific, Waltham, MA, USA). Reactions were performed using a Rotor-Gene 6000 cycler (Corbett Research, Sydney, NSW, Australia). A standard curve was obtained using dilutions of the cDNA sample. Expression was quantified using ROTOR-GENE software (Corbett Research). Four independent technical repeats were performed for each sample. Relative expression was calculated by dividing the expression level of the examined gene by that of *SlACTIN*. The target-gene-to-*ACTIN* ratio was then averaged. The values for control treatments were set to 1. All primer sequences are presented in Supplementary Table S1.

### Library Constructions and RNAseq analysis

Total RNA was extracted from terminal leaflets of the third fully expanded leaf using a RNeasy Plant Mini Kit (Qiagen, Hilden, Germany). RNA-seq libraries were prepared at the Crown Genomics Institute of the Nancy and Stephen Grand Israel National Center for Personalized Medicine (INCPM), Weizmann Institute of Science, Israel, using the INCPM-mRNA-seq library preparation protocol. Briefly, the poly(A) fraction (mRNA) was purified from 500 ng of total input RNA, followed by fragmentation and synthesis of double-stranded cDNA. Libraries were purified using Agencourt AMPure XP beads (Beckman Coulter Life Sciences, Indianapolis, IN, USA), and subjected to end repair, A-tailing, adapter ligation, and PCR amplification. Library quality and concentration were assessed using a Qubit Fluorometer (Thermo Fisher Scientific, Waltham, MA 15 USA) and a TapeStation system (Agilent Technologies, Santa Clara, CA, USA). Sequencing was performed on an Illumina NextSeq platform (Illumina, San Diego, CA, USA) using a 75-cycle high output kit, generating 75 bp single-end reads. Approximately 39 million reads were obtained per sample.

### Sequence data analysis

Poly-A/T stretches and Illumina adapters were trimmed from the reads using cutadapt (Martin, 2011); resulting reads shorter than 30bp were discarded. Reads were mapped to the reference genome *Solanum lycopersicum* release SL4.0 using STAR (Dobin *et al.,* 2013), (with End to End option and out Filter Mismatch Nover Lmax set to 0.04). Annotation file was downloaded from SolGenomics, ITAG release 4.0. Reads with the same UMI were removed using the PICARD MarkDuplicate tool using the BARCODE_TAG parameter. Expression levels for each gene were quantified using htseq-count (Anders *et al.,* 2015), using the GTF (Gene Transfer Format) file downloaded from SolGenomics. Differentially expression analysis was performed using DESeq2 (Love *et al.,* 2014) with the betaPrior, cooksCutoff and independent filtering parameters set to False. Raw P values were adjusted for multiple testing using the procedure of Benjamini and Hochberg (1995). Pipeline was run using snakemake (Köster and Rahmann, 2012).

### Microscopy and Image Analysis

#### Hypocotyl Imaging

Hypocotyl tissues from seedlings were divided into three regions: upper, middle, and bottom. Hand-cut cross-sections from each region were mounted vertically on metal stubs coated with carbon tape. Imaging was performed using a Phenom ProX Scanning Electron Microscope (SEM) (Thermo Fisher Scientific, Waltham, MA, USA) equipped with a Backscatter Electron Detector (BSD). Images were acquired and processed using Phenom ProSuite software. Epidermal cell length was measured using ImageJ software (https://imagej.net/ij/). For each genotype, the final reported value represented the average epidermal cell length across the upper, middle, and bottom regions.

#### Embryo Imaging

To isolate embryos, tomato seeds were imbibed in water for several hours and then dissected using a surgical blade and fine tweezers under a binocular microscope (Olympus, Waltham, MA, USA). Rescued embryos were imaged using a Nikon SMZ1270 stereo microscope equipped with a Nikon DS-Ri2 camera and NIS-Elements software (Nikon Instruments, Melville, NY, USA). Embryo and cotyledon lengths were measured using ImageJ software.

#### Hypocotyl Diameter Imaging (Light Microscopy)

Hand-cut hypocotyl cross-sections from seedlings were captured using a Leica DM500 microscope equipped with a Leica ICC50 W camera and Leica Application Suite software (Leica Microsystems, Heerbrugg, Switzerland). Hypocotyl diameters were analyzed using ImageJ software.

### Statistical analysis

All assays were conducted with three or more biological replicates and analyzed using JMP software (SAS Institute, Cary, NC, USA). Means comparison was conducted using analysis of variance (ANOVA) followed by post-hoc tests. For multiple comparisons among all groups, the Tukey–Kramer honest significant difference (HSD) test was used. For comparisons of experimental groups specifically against control, Dunnett’s test was applied. For single comparisons between two groups, Student’s *t*-test was used. A significance threshold of *P* < 0.05 was applied for all tests.

### Gene annotation and accession numbers

Sequence data from this article can be found in the Sol Genomics Network (https://solgenomics.net/) under the following accession numbers: *ACTIN, Solyc11g005330; GA2ox1, Solyc05g053340; GA2ox2, Solyc07g056670; GA2ox4,Solyc07g061720; GA2ox7, Solyc02g080120, GA20ox1, Solyc03g006880; GA20ox2, Solyc06g035530; GA20ox3, Solyc11g072310; GA20ox4, Solyc01g093980; GA3ox1, Solyc06g066820; GA3ox2, Solyc03g119910; GA3ox3, Solyc01g058250; GA3ox4, Solyc05g052740; GA3ox5, Solyc00g007180; GID1b1, Solyc09g074270; TINY1, Solyc06g066540; TINY2, Solyc03g120840; TINY3 ,Solyc12g044390; TINY4, Solyc12g008350; TINY5, Solyc08g066660; TINY6, Solyc01g090560; Lectin-domain receptor-like kinas, Solyc07g055690; SEC3A, Solyc07g025170; SIP5CS1, Solyc06g019170; RD29A, Solyc03g025810; RD29B, Solyc01g009660; SlRAB18, Solyc02g084850; PROCERA, Solyc11g011260*.

## Results

### The loss of TINY1 increased the rate of water loss under drought conditions

Previously, we demonstrated that the tomato *TINY1* is induced by drought. In Arabidopsis, TINY1 regulates stomatal closure and influences the rate of water loss (Xie *et al*., 2019). To assess whether TINY1 activity in tomato also affects water loss, we grew WT M82 and *tiny1* mutant plants under normal irrigation conditions for five weeks after which irrigation was stopped. We then monitored the rate of water loss. *tiny1* plants lost their turgor faster than the M82 (7 days on average into the drought treatment), and the relative water content (RWC) of the leaves at that time was significantly lower in the mutant (Fig. 1A and B).

**Figure 1.**
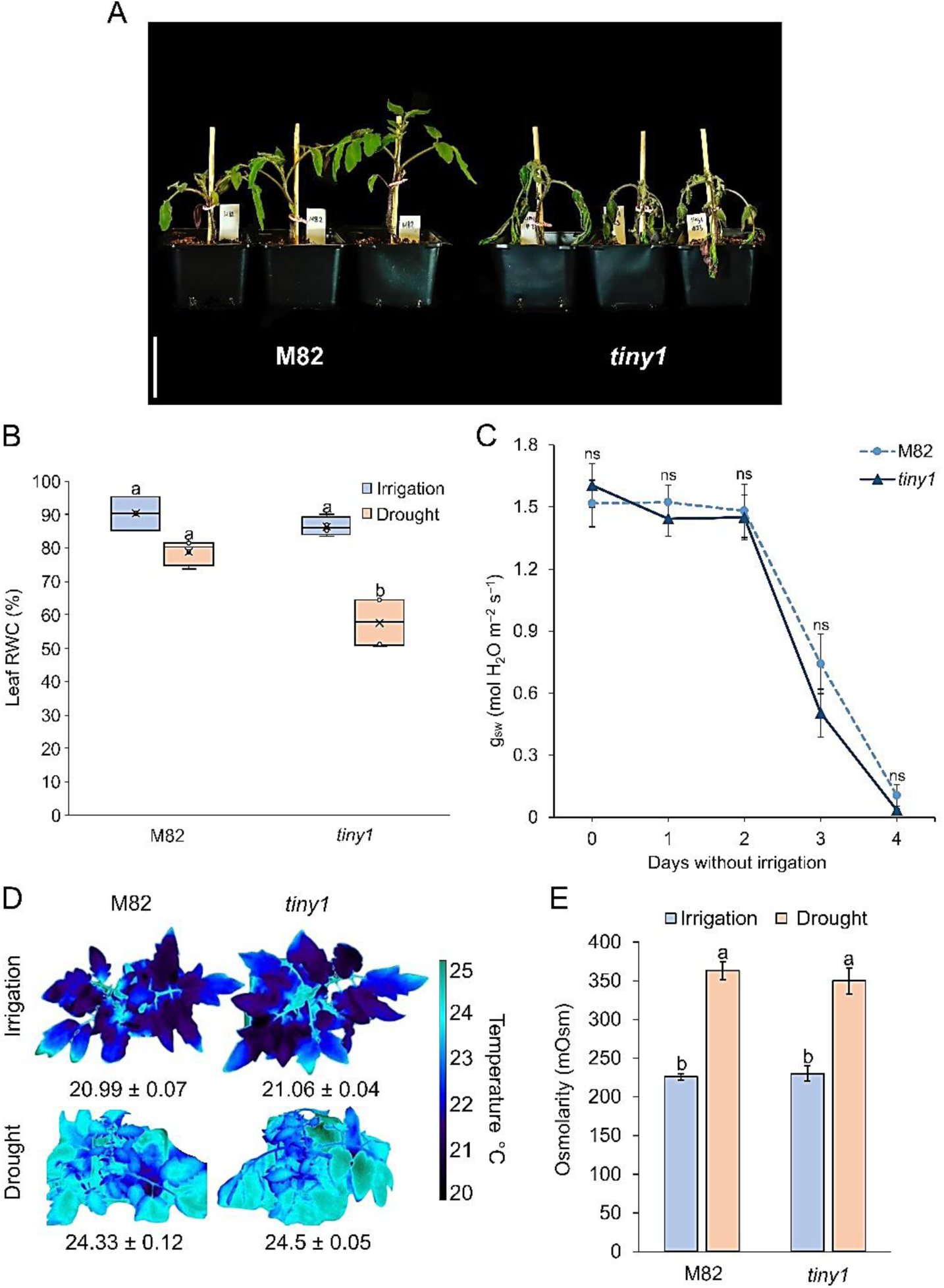
*tiny1* exhibited faster water loss under drought conditions. **A.** Representative M82 and *tiny1-23* plants after eight days without irrigation. Scale bar= 6 cm. **B**. Leaf relative water content (RWC, %) of M82 and *tiny1-23* under normal irrigation and seven days without irrigation. Values are means± SE of six biological replicates (terminal leaflets taken from the third leaf below the apex). Different letters above box plots indicate statistically significant differences (P < 0.05, Tukey-Kramer HSD). **C**. Stomatal conductance (g_sw_) of M82 and *tiny1-23*, measured at 9:00 AM over four days without irrigation. Data represent means± SE of 14 biological replicates for M82 and 19 biological replicates for *tiny1-23*, with three technical replicates per measurement. Statistical significance was determined using Student’s *t*-test (P < 0.05); “ns“ denotes non-significant differences. D. Thermal imaging of representative M82 and *tiny1-23* plants at 10:00 AM under irrigation and three days after irrigation was stopped. Numbers below plants are the means± SE leaf-surface temperature taken from the entire documented plants and the values are means of 9 biological replicates. **E.** Osmolarity of terminal leaflets from the third fully expanded leaf below the apex from M82 and tiny1-23 plants grown under irrigation or exposed to drought conditions (85% leaf RWC for irrigation and 65% leaf RWC for drought). Data are the means± SE of five biological replicates per genotype. Different letters above the plots indicate statistically significant differences (P < 0.05, Tukey-Kramer HSD).

We then examined stomatal conductance in irrigated plants and during soil dehydration. We did not observe any differences in stomatal conductance between the WT and *tiny1* under either well-watered or drought conditions (Fig. 1C). Transpiration was further assessed using thermal imaging, since evaporative cooling via transpiration is a major driver of leaf temperature regulation. Both genotypes exhibited similar leaf temperatures during soil dehydration, indicating comparable transpiration rates (Fig. 1D). Next, we tested the effect of drought on osmolyte accumulation in the leaves of M82 and *tiny1* plants, both under well-watered conditions and 7 days after irrigation was withheld. No differences were observed between the two genotypes (Fig. 1E). These results do not support the hypothesis that the faster water loss found in *tiny1* is due to changes in stomatal conductance and transpiration rate.

### The increased rate of water loss in *tiny1* is due to larger canopy area

Larger leaf area leads to increased total transpiration, faster soil dehydration, and earlier loss of turgor. Since it was shown previously that TINY suppresses growth (Xie *et al*., 2019), we compared the leaf area (transpirational surface area) between WT M82 and *tiny1*. M82 and *tiny1* were grown for six weeks and then total leaf area and the number of leaves were measured. The foliage area of *tiny1* was larger than that of M82 (Fig. 2A), and the mutant had, on average, one more leaf than M82 (Fig. 2B). To determine if this causes faster water loss and faster wilting of *tiny1*, we grew the mutant and M82 plants side by side in the same pot. This approach eliminates the effect of plant size on the time of turgor loss, since both genotypes are exposed to the same volumetric soil water content throughout the experiment (Shohat *et al*., 2021). After six weeks of growth, irrigation was stopped. Six days into the drought treatment, both M82 and *tiny1* plants lost turgor simultaneously and began wilting (Fig. 2C). At this time point, leaf RWC was similar in the two lines (Fig. 2D). We also analyzed whole-plant transpiration in M82 and *tiny1* plants, grown in a greenhouse using an array of lysimeters (Illouz Eliaz *et al*., 2020). Since there is some degree of variation in plant size (leaf number) in the two genotypes, we were able to select for plants with similar number of leaves. Plants with 7 leaves (M82 and *tiny1*) were grown for 10 days on the lysimeters with irrigation, and then divided into three groups: group 1 continued to receive normal irrigation, group 2 were exposed to moderate drought (received 50% irrigation of their daily transpiration) and for group 3 water supply was terminated. Whole plant transpiration rate, three days after the beginning of the treatments, was similar between M82 and *tiny1* under all tested conditions (Fig. 2E). Together, these results suggest that the accelerated wilting of *tiny1* is likely due to its larger leaf area, rather than any inherent differences in dehydration response.

**Figure 2.**
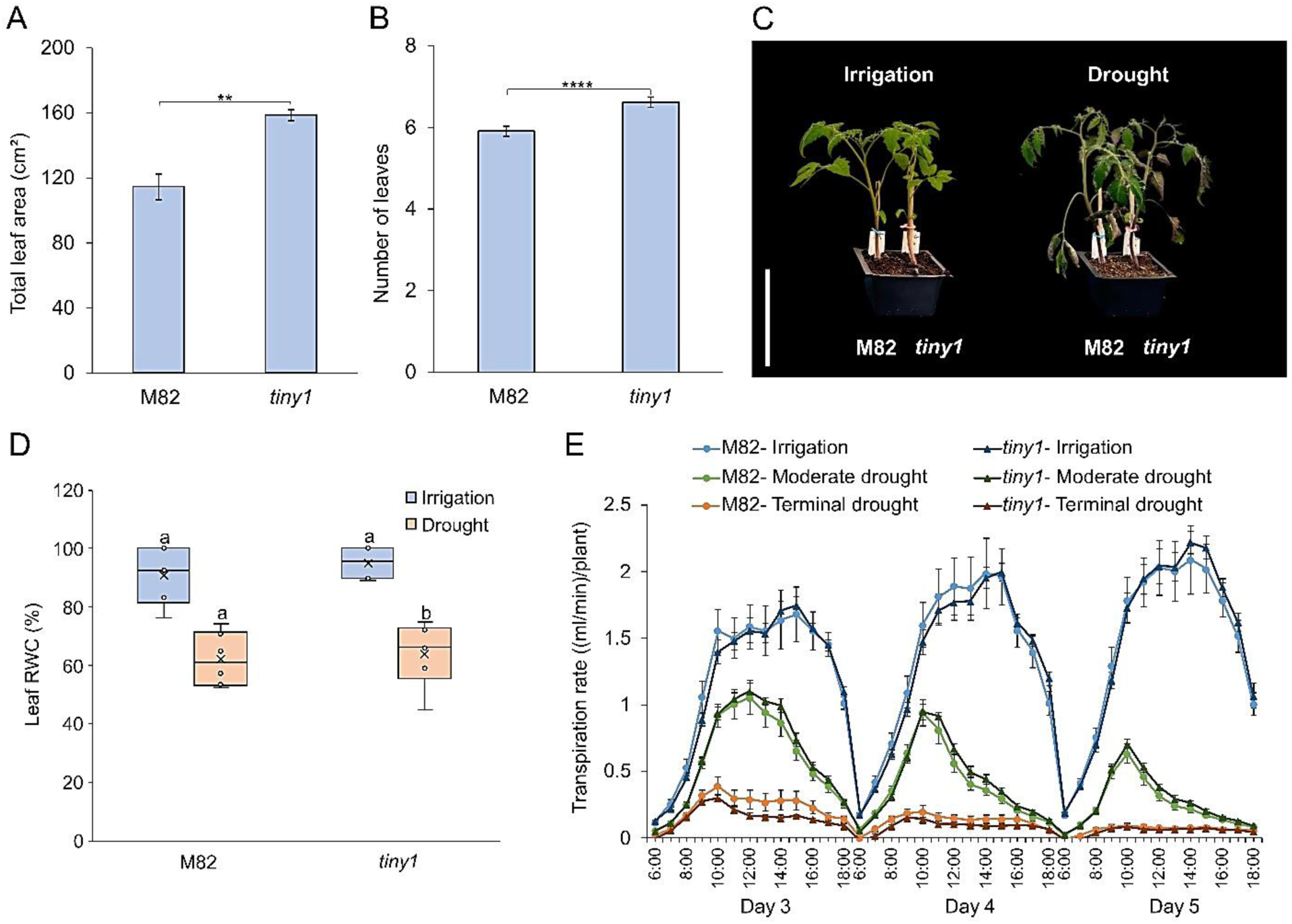
The rapid water loss of *tiny1* under drought is due to its larger leaf area. **A.** Total leaf area (cm²) of five-week-old M82 and *tiny1-23* plants. Values are mean ± SE of five plants. **B**. Number of leaves in five-week-old M82 and *tiny1-23* plants. Values are the means± SE of 40 plants. Asterisks in panels A and B represent significant differences between genotypes by Student’s t-test (P < 0.05). **C**. M82 and tiny1-23 plants were grown side by side in the same pot and after six weeks irrigation was stopped. Images show representative M82 *and tiny1-23* plants under irrigation and five days after irrigation was withheld. Scale bar= 15 cm. **D**. Leaf RWC (%) of M82) and *tiny1-23* under irrigation and drought conditions (five days without irrigation). Values are means± SE of six biological replicates (terminal leaf taken from the third leaf below the apex). Different letters above box plots indicate statistically significant differences (P < 0.05, Tukey-Kramer HSD). E. Whole-plant transpiration rates of M82 and *tiny1-23* under different irrigation regimes. Transpiration was measured continuously over 72 hours (06:00–18:00) during days 3, 4, and 5 of the experiment under three irrigation conditions: Normal irrigation (control), moderate drought (irrigation of 50% of daily transpiration) and terminal drought (no irrigation). Values represent means ± SE for seven plants per genotype in the control, nine plants per genotype in the moderate drought, and eight plants per genotype in the terminal drought condition. M82 and *tiny1-*23 plants with the same number of leaves (7 expanded leaves) were placed on lysimeters, and pot weight (pot + soil + plant) was recorded every three minutes.

### The effect of TINY1 on global transcriptional activity under drought conditions

We next explored the effect of *tiny1* on leaf transcriptional response under drought by performing RNAseq analysis. M82 and *tiny1* plants were grown for five weeks under normal irrigation conditions, after which irrigation was stopped. Once the plants lost turgor (RWC 50%), terminal leaflets from leaf number 3 (top down) were collected for RNA extraction and RNAseq analysis. Using a 2-fold increase or decrease cutoff (adjusted P value for multiple comparisons ≤0.05) and filter out low read count (<30 reads), we identified 4545 differentially expressed genes (DEGs) between irrigated and dehydrated M82 plants (1438 upregulated and 3107 downregulated) and 3610 DEG in *tiny1* plants (1198 upregulated and 2412 downregulated) (Fig. 3A and Supplementary Dataset S1). Most of the DEGs found in M82 were also DEG in *tiny1*. We found 71 DEG that were upregulated in M82 under drought conditions but were not affected or significantly less affected in *tiny1* (Supplementary Dataset S2). The most significant DEG induced by drought in M82 but not affected in *tiny1* was *LECTIN-DOMAIN RECEPTOR-LIKE KINASE*, Solyc07g055690 (Fig. 3B). Among the downregulated DEGs we did not find clear putative targets of TINY1. Regardless drought conditions, we identified one clear putative target of TINY1, *EXOCYT COMPLEX COMPONENT 3A*, Solyc07g025170 (SEC3A). The expression of this gene was slightly but not significantly reduced under drought conditions in M82, but in *tiny1* it was not expressed under irrigation or drought conditions (Fig. 3C). We confirm these results by RTqPCR (Supplementary Fig. S2). We also evaluated the effect of drought on known drought regulated genes *RD29a-*Solyc03g025810; *RD29B*-Solyc01g009660, *SlP5CS1*-Solyc06g019170; and *SlRAB18*-Solyc02g084850 (Nir *et al*., 2017). While the expression of *RD29A* was slightly lower under drought in *tiny1*, the other genes were similarly affected by drought in M82 and *tiny1* (Fig. 3D). These results suggest that the loss of TINY1 activity has minor, effect on the leaf transcriptional response to drought. We also examined the expression of the different TINY genes in M82. *TINY1*, *TINY2*-Solyc03g120840 and *TINY3*-Solyc12g044390 were strongly upregulated by drought, whereas *TINY4*-Solyc12g008350 and *TINY5*-Solyc08g066660 were downregulated by drought (Fig. 3E). The expression of *TINY6*-Solyc01g090560 was not affected. It is possible that the loss of TINY1 activity did not affect plant responses to drought (stomatal activity and gene expression) due to functional redundancy with TINY2 and TINY3.

**Figure 3.**
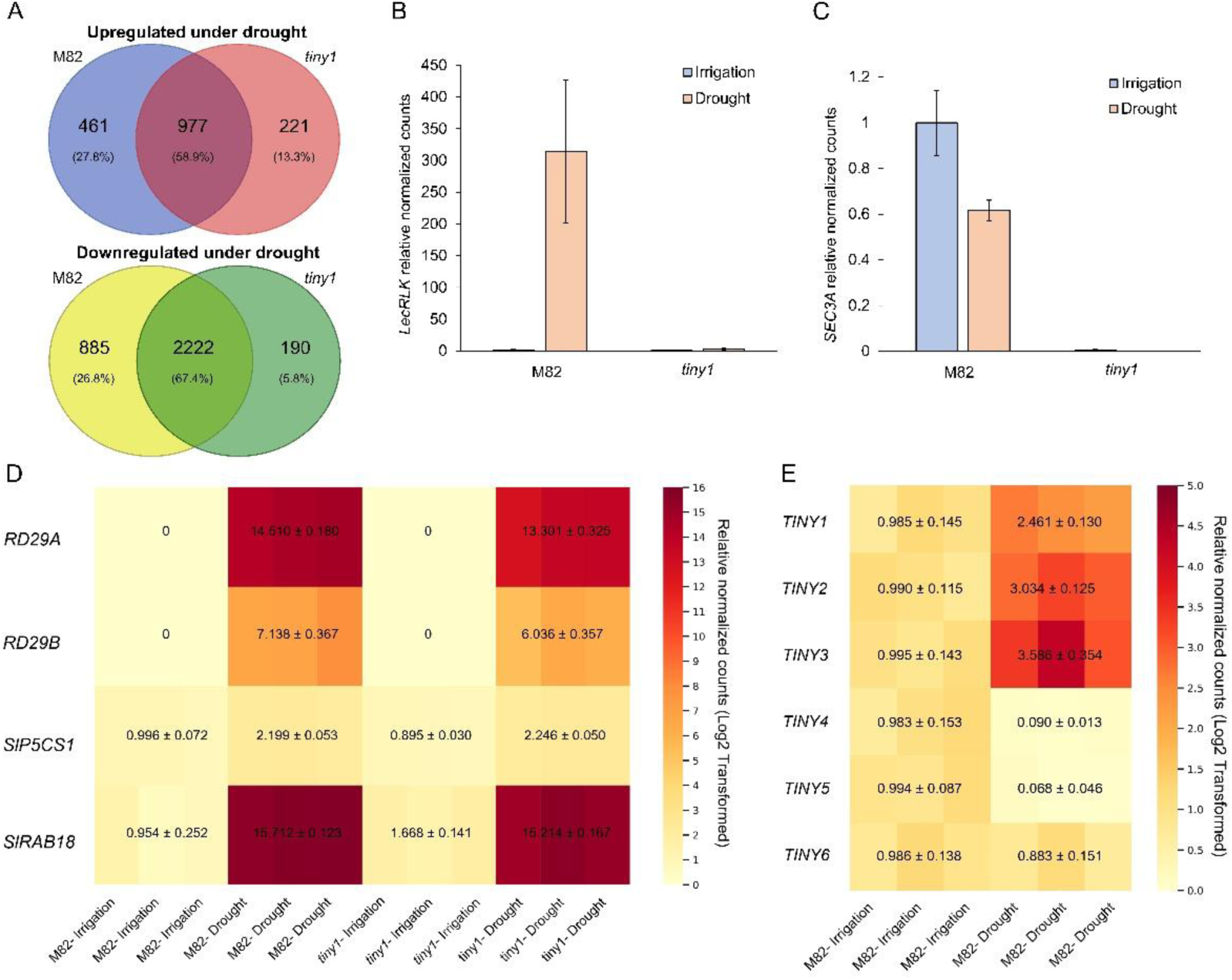
Global transcriptional drought response in *tiny1* leaves. **A.** Venn diagrams showing the overlap of differentially expressed genes (DEG) between drought (51-55% leaf RWC) and irrigation (80-85% leaf RWC) in M82 and *tiny1-23*. DESeq2 analysis was performed on RNA-Seq data to identify upregulated genes (top panel) and downregulated genes (bottom panel). Numbers indicate the number of DEG, with percentages representing their proportion of the total DEG. **B**. Relative normalized counts of *LECTIN-DOMAIN RECEPTOR-LIKE KINASE* (*LecRLK*) in M82 and *tiny1-23* leaves under irrigation (80-85% leaf RWC) and drought (51-55% leaf RWC). **C**. Relative normalized counts of *EXOCYST COMPLEX COMPONENT 3A* (*SEC3A*) in M82 and *tiny1-23* leaves under irrigation (80-85% leaf RWC) and drought (51-55% leaf RWC). (B and C) Values are means ±SE of three biological replicates. **D**. Heatmap of log₂-transformed relative normalized expression levels of drought-responsive genes under irrigation (80-85% leaf RWC) and drought (51-55% leaf RWC) in M82 and *tiny1-23*. Each treatment includes three biological replicates. Drought-induced genes with their respective Solyc IDs from the Sol Genomics Network are indicated. **E**. Heatmap of log₂-transformed relative normalized expression levels of TINY genes in M82 under irrigation (80-85% leaf RWC) and drought (51-55% leaf RWC). TINY genes with its Solyc ID from the Sol Genomics Network are indicated. The color gradient in D and E represents expression levels, with light yellow indicating low expression and dark red representing high expression. Values are presented as means± SE of log₂ (normalized expression + 1). The scale bar on the right denotes expression intensity.

### TINY1 suppressed embryonic and early seedling development

To find at what developmental stage the loss of TINY1 activity affects growth, we followed the rate of leaf production and found that *tiny1* developed the first true leaf earlier than the M82, but later the rate of leaf production was similar Supplementary Fig. S3). These results suggest that the effect of TINY1 on growth is restricted to early stages of plant development. We found that the seeds of *tiny1* (lines *tiny1-1* and *tiny1-23*) were larger than those of M82 (Fig. 4A and Supplementary Fig. S4). We therefore extracted the embryos from the seeds and measured their size. The embryos of *tiny1* were much larger than those of the M82 (Fig. 4B and Supplementary Fig. S5). We also examined the seedlings and found that two weeks after germination the hypocotyl of *tiny1* was longer than in M82 (Fig. 4C and Supplementary Fig. S6). Microscopic analyses of the hypocotyl’s epidermal cells and cross sections of the hypocotyl suggest the TINY1 affect cell elongation and expansion rather than cell division (Fig. 4D and E and Supplementary Fig. S7S and B). Together, these results suggests that TINY1 act as a growth suppressor during the early stages of plant development.

**Figure 4.**
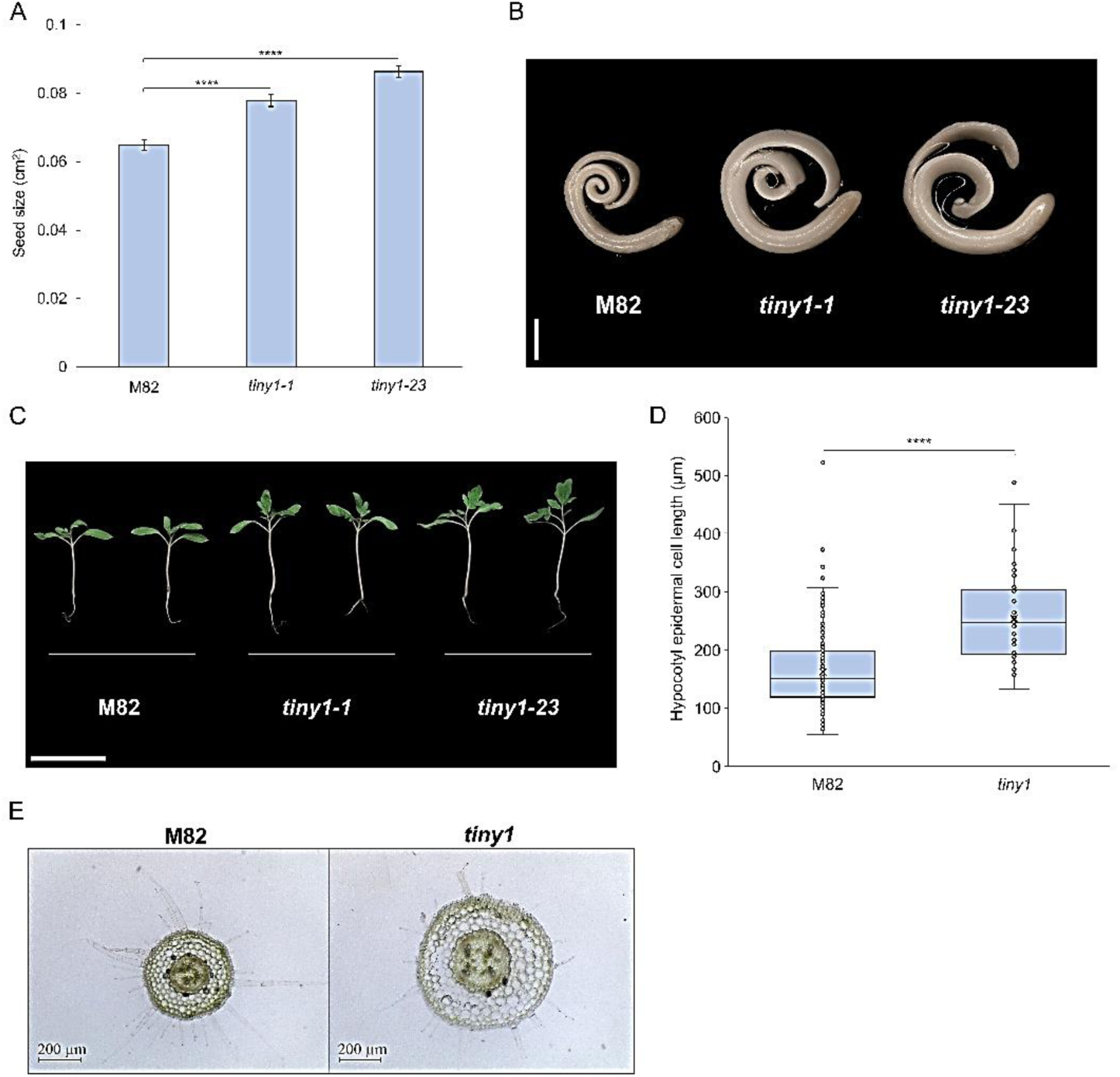
TINY1 suppressed embryo and seedling growth. **A.** M82, *tiny1-1* and *tiny1-23* seed size (cm^2^). Values are means ± SE of 30 seeds. Asterisks indicate significant differences between genotypes by Dunnett test (P < 0.05). **B**. Embryos of M82, *tiny1-1* and *tiny1-23* extracted from dry seeds. Scale bar = 1 mm. **C**. Representative seedlings of M82, *tiny1-1* and tiny*1-23* 18 days after germination. Scale bar= 5 cm. **D**. Hypocotyl epidermal cell length measured in two-week-old seedlings of M82 and *tiny1-23*, Values are means ± SE of 230 cells of M82 and 300 cells of *tiny1* . Asterisks represent significant differences between genotypes by Student’s t-test (P < 0.05). **E**. Representative cross-sections of hypocotyls from twenty-day-old M82 and *tiny1-23* seedlings. Scale bar=200 µm.

TINY1 expression is induced by drought. Since we found evidence that TINY1 acts as embryonic growth suppressor, we hypothesized that its role in drought conditions is to suppress plant growth. We therefore grew M82 and *tiny1* plants under mild drought conditions that reduce growth rates. The plants were grown for three weeks under regular irrigation, after which irrigation was stopped. When the plants began to lose turgor, 40% of the normal irrigation volume was added. This treatment was repeated for five cycles, lasting approximately 35 days. At the beginning of the drought treatment and just before each subsequent irrigation, stem length was measured and leaf number counted. At the end of the experiment, leaf area was measured. Drought treatment inhibited stem elongation and leaf production, but the effect was similar in M82 and *tiny1* (Supplementary Fig. S8A and B). The total leaf area was strongly reduced by drought, but again, the loss of TINY1 activity did not affect this growth rate (Supplementary Fig. S8C). This suggests that either TINY1 has no role in regulating plant growth under water-limited conditions, or its function is completely redundant with the activity of other TINY proteins.

### TINY1 attenuated GA activity in the maturing embryo

Previous studies in tomato suggest that TINY1 regulates GA biosynthesis in leaves via the inhibition of *GA20ox1*, *GA20ox2* and *GA20ox4* expression (Li *et al*., 2012) and GA catabolism in guard cells by the induction of *GA2ox7* (Shohat et al., 2021). We therefore generated *tiny1* homozygous plants on the *PROCERA* (*pro*)*Δ17* background (Nir *et al*., 2017). *proΔ17* plants overexpress a gain-of-function stable version of the DELLA protein PRO that constitutively inhibits GA activity. The double mutant *tiny1 proΔ17* seedlings exhibited the dwarf phenotype (short hypocotyl) of *proΔ17* and leaf number were as in M82 and *proΔ17* but less than *tiny1* (Fig. 5A, B and Supplementary Fig. S9), suggesting that PRO is epistatic to TINY1 and support the hypothesis that TINY1 regulates growth via its effect on GA biosynthesis/catabolism. To test if GA activity increases in *tiny1* mutant, we analyzed the expression of *GID1b1* (GA receptor gene). *GID1b* expression is inhibited by increased GA activity due to a feedback response (Illouz-Eliaz *et al*., 2019). The expression of *GID1b1* was analyzed in seeds extracted from red fruits, in seedlings following germination and in elongating hypocotyls two weeks after germination. *GID1b1* expression was inhibited in *tiny1* (compared to M82) seeds but not in seedlings or in elongating hypocotyl (Fig. 5C), suggesting increased GA activity in the mutant embryos, but not later during seedling development. We then analyzed the candidate targets of TINY1: *GA20ox1*, *GA20ox2*, *GA20ox4* and *GA2ox7* in seeds extracted from red fruits. The expression of *GA20ox4*, but not of the other examined genes was significantly affected (upregulated) by *tiny1* (Fig. 5D-G). We further analyzed other *GA20ox*, *GA3ox* and *GA2ox* genes but none of them was affected significantly by the mutation (Supplementary Fig. S10). These results suggest that TINY1 suppresses the expression of *GA20ox4* in the embryo to reduce GA production.

**Figure 5.**
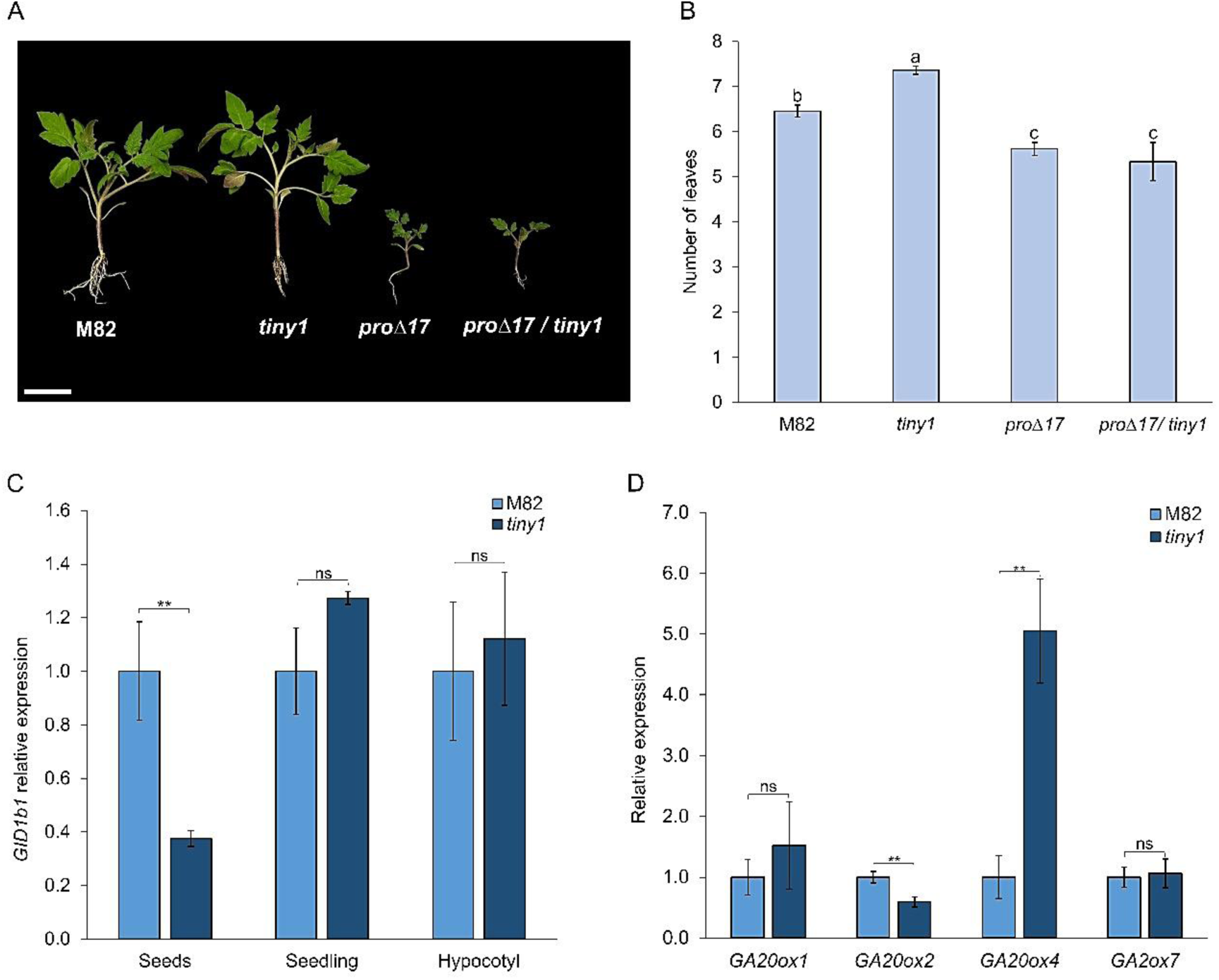
TINY1 repressed embryo growth via the GA pathway. **A.** Representative seedlings of M82, *tiny1-23*, 35S:proΔ17, and 35S:proΔ17/*tiny1-*23. **B.** Number of leaves in 30-day-old M82, *tiny1-23*, 35S:proΔ17, and 35S:proΔ17/*tiny1-23* plants. Data are means ± SE of 37 (M82), 33 (*tiny1-23*), 13 (35S:proΔ17), and 6 (35S:proΔ17/*tiny1-23*) plants. Different letters indicate statistically significant differences between lines (P < 0.05, Tukey-Kramer HSD test). **C**. Relative expression (RT-qPCR) of *GID1b1* in seeds harvested from red-ripe fruit, in one week old seedlings and in hypocotyls of M82 and *tiny1-23*. Expression levels are normalized to M82 seeds and presented as means ± SE of minimum 4 replicates. **D**. Relative expression (RT-qPCR) of GA biosynthesis and deactivation genes (*GA20ox1*, *GA20ox2*, *GA20ox4* and *GA20ox7*) in seeds harvested from red-ripe fruit of M82 and *tiny1-23*. Expression levels are normalized to M82 and values are means ± SE of 4 biological replicates. Asterisks in C and D indicate significant differences between genotypes (P < 0.05, Student’s t-test); “ns“ denotes no significant difference.

## Discussion

The transcription factors DREB play an important role in plant responses to various abiotic and biotic stresses, including drought. Many studies have shown their effect on drought avoidance responses (stomatal closure and growth suppression under drought) and drought tolerance (regulation of drought-responsive genes) (Lata and Prasad, 2011; Sarkar *et al*., 2019). This family of proteins is divided into six subfamilies (A1 to A6). A subgroup of subfamily A4 is named TINY (Nakano *et al*., 2006). TINY proteins seem to regulate both growth and abiotic responses. TINY1 overexpressing Arabidopsis and tomato plants are semi-dwarf (Wilson *et al*., 1996; Li *et a*l., 2012). Moreover, the triple Arabidopsis mutant *tiny1tiny2tiny3* has larger leaves and longer petioles (Xie *et al*., 2019). In both plants overexpression of TINY1 improved drought tolerance due to increased drought-responsive gene expression and hypersensitivity to ABA-mediated stomatal closure (Sun *et al*., 2008; Xie *et al*., 2019).

We found that the loss of TINY1 gene in tomato resulted in early wilting when exposed to water limited conditions (Fig. 1A-B). However, this rapid water loss was not associated with increased stomatal conductance or impaired osmotic adjustment (Fig. 1C-E); but rather with an increased leaf area which led to greater total transpirational surface (Fig. 2A-B). *tiny1* mutant produced its first true leaf earlier than the control M82, resulting in the development of, on average, one additional leaf during early vegetative growth before the onset of its first inflorescence (Fig. S2). RNAseq analysis of leaves taken from well-watered and drought treated M82 and *tiny1* plants, revealed similar expression of known drought responsive genes (Fig. 3D). This however does not necessarily suggest that TINY proteins have no effect on drought-related gene expression. Three *TINY* genes were upregulated by drought in leaves, *TINY1*, *TINY2* and *TINY3*, it is therefore possible that *TINY2/3* compensated for the loss of *TINY1* in the regulation of drought-induced molecular responses in leaves (Fig. 3E), due to functional redundancy. We identified, however, a potential drought-induced TINY1 target: *LECTIN-DOMAIN RECEPTOR-LIKE KINASE*, whose expression was strongly upregulated by drought in M82 leaves but not in *tiny1* (Fig. 3B). Members of this receptor family have been shown to play key roles in plant development as well as in responses to both abiotic and biotic stresses (Vaid *et al*., 2013; Sun et al., 2020). Additionally, our analysis revealed the tomato exocyst subunit *SEC3A* as another potential TINY1 target in leaves (Fig. 3C). Although *SEC3A* expression was not affected by drought, it was completely suppressed in all *tiny1* samples, regardless of water availability. Previous studies have implicated SEC3A in several key processes, including cell growth (Hála *et al*., 2008), embryo development (Zhang et al., 2013), and pathogen defense (Du *et al*., 2018). Based on these findings, the loss of exocyst activity would be expected to result in smaller embryo and shorter hypocotyls. However, *tiny1* mutants exhibited larger embryo and longer hypocotyls, thus, the developmental role of TINY1 in regulating *SEC3A* remains unclear.

The only clear and consistent phenotypes we observed in *tiny1* were larger seeds and embryos, longer hypocotyls, and early emergence of the first true leaf (Fig. 4A-D). At later stages of seedling and plant development we did not find differences in plant growth rate and development. As mentioned above, the effect of TINY proteins on plant growth was shown before (Wilson *et al*., 1996; Li *et al*., 2012; Xie *et al*., 2019). Here we show that in tomato its effect on plant size resulted from its regulation of embryo and early seedling development. Xie *et al*. (2019) proposed that the Arabidopsis TINY proteins are regulated by BR and, in turn, modulate BR signaling. Under drought conditions, the kinase BIN2 phosphorylates and stabilizes TINY proteins. The accumulated TINY then interacts with and inhibits the BR-responsive transcription factor BES1, thereby repressing BR-mediated growth and activating drought-responsive genes. In contrast, under normal conditions, active BR signaling promotes the degradation of TINY proteins, which promotes growth and suppresses stress-related gene expression. In tomato on the other hand, TINY1 overexpression suppressed GA biosynthesis (Li *et al*., 2012), suggesting that the effect of TINY on growth is via the GA response pathway. Our results also suggest that TINY1 promotes embryo growth through the GA pathway. Loss of TINY1 function strongly induced the expression of the GA biosynthesis gene *GA20ox4* in maturing seeds (Fig. 5D). Elevated GA activity typically triggers a feedback response that, in tomato, includes suppression of the GA receptor gene *GID1b1* (Illouz-Eliaz *et al*., 2019). The reduced expression of *GID1b1* in *tiny1* maturing seeds further supports the notion of increased GA activity (Fig. 5C). In addition, the excess hypocotyl elongation and the early leaf growth phenotype observed in *tiny1* were suppressed in the presence of a stabilized DELLA protein (Fig. 5A–B), suggesting that TINY1 acts upstream of DELLA in the GA pathway. Although DELLA proteins also influence the BR pathway (Shani *et al*., 2024), our collective findings indicate that in developing embryos, TINY1 primarily inhibits the GA pathway to suppress growth. While *tiny1* mutants displayed enhanced embryo size, longer hypocotyls, and accelerated emergence of the first true leaves, we detected altered GA activity only in the maturing embryo. This may suggest that GA induces stable epigenomic changes in growth-related genes that persist following seed dormancy and germination. Alternatively, TINY1 may act via different pathways to regulate embryo growth and hypocotyl elongation, e.g. in the embryo it suppresses GA activity and, in the hypocotyl, BR activity.

To conclude, the tomato TINY1 probably has two different functions; an embryonic growth suppressor and, a yet unknown function in leaves under drought. The inhibition of embryo growth and hypocotyl elongation by TINY1 may contribute to improved seedling resilience following germination under changing environments.

## Acknowledgements

We would like to thank Dr. Naama Teboul and Dr. Adi Yaaran for technical assistance.

## Author Contributions

MA, YZ and DW designed the research; MA, HS and DB performed experiments; MA, DW and YZ analyzed data; MA, YZ and DW wrote the manuscript.

## Data Availability

All data can be found in the manuscript and in the online Supplementary Data. Sequence data (RNAseq) from this article can be found in the Sol Genomics Network (https://solgenomics.net/)

## Supplementary material

The following supplementary material are available.

**Supplementary Figure S1.** Sequence analysis of the *tiny1-23* allele.

**Supplementary Figure S2**. *EXOCYT COMPLEX COMPONENT 3A* (SEC3A) expression in M82 and *tiny1.*

**Supplementary Figure S3.** Rate of leaf production in M82 and *tiny1-23* plants.

**Supplementary Figure S4.** Seeds of M82 and *tiny1* mutants.

**Supplementary Figure S5.** M82 and *tiny1* embryo size.

**Supplementary Figure S6.** M82 and *tiny1* hypocotyl length.

**Supplementary Figure S7.** The loss of TINY1 activity promoted hypocotyl cell elongation and expansion.

**Supplementary Figure S8.** The loss of TINY1 activity did not promote plant growth under drought.

**Supplementary Figure S9.** Stable DELLA protein inhibited the effect of tiny1 on seedling growth.

**Supplementary Figure S10.** Expression of GA biosynthesis and catabolism genes in seeds of M82 and *tiny1*.

**Supplementary Table S1.** Primers used in this study.

**Supplementary Dataset S1.** Differentially expressed genes (DEG) between irrigation and drought conditions in M82 and *tiny1*.

**Supplementary Dataset S2.** Differentially expressed drought-induced genes between M82 and *tiny1*.

## Abbreviations

DEG: differentially expressed gene
DREB: Dehydration-Responsive Element-Binding
RWC: relative water content

## References

Agarwal PK, Agarwal P, Reddy MK, Sopory SK. 2006. Role of DREB transcription factors in abiotic and biotic stress tolerance in plants. Plant Cell Reports 25, 1263–1274.

Ali O, Cheddadi I, Landrein B, Long Y. 2023. Revisiting the relationship between turgor pressure and plant cell growth. New Phytologist 238, 62–69.

Anders S, Pyl PT, Huber W. 2015. HTSeq- a Python framework to work with high-throughput sequencing data. Bioinformatics. 31, 166–169. 10.1093/bioinformatics/btu638

Benjamini Y, Hochberg Y. 1995. Controlling the false discovery rate: a practical and powerful approach to multiple testing. Journal of the Royal Statistical Society Series B 57, 289–300.

Bharath P, Gahir S, Raghavendra AS. 2021. Abscisic acid-induced stomatal closure: An important component of plant defense against abiotic and biotic stress. Frontiers in Plant Science 12, 615114. 10.3389/fpls.2021.615114

Blum A. 2017. Osmotic adjustment is a prime drought stress adaptive engine in support of plant production. Plant Cell and Environment 40, 4–10.

Dalal A, Shenhar I, Bourstein R, Mayo A, Grunwald Y, Averbuch N, Attia Z, Wallach R, Moshelion M. 2020. A telemetric, gravimetric platform for real-time physiological phenotyping of plant–environment interactions. Journal of Visualized Experiments 162, 61280. 10.3791/61280

Dobin A, Davis CA, Schlesinger F, Drenkow J, Zaleski C, Jha S, Batut P, Chaisson M, Gingeras TR. 2013. STAR: ultrafast universal RNA-seq aligner. Bioinformatics 29, 15–21. 10.1093/bioinformatics/bts635

Du Y, Overdijk EJR, Berg JA, Govers F, Bouwmeester K. 2018. Solanaceous exocyst subunits are involved in immunity to diverse plant pathogens. Journal of Experimental Botany 69, 655–666.

Edgar R, Domrachev M, Lash AE. 2002. Gene Expression Omnibus: NCBI gene expression and hybridization array data repository. Nucleic Acids Research 30, 207–210.

Gupta A, Rico-Medina A, Caño-Delgado AI. 2020. The physiology of plant responses to drought. Science 368, 266–269.

Hála M, Cole R, Synek L, et al. 2008. An exocyst complex functions in plant cell growth in *Arabidopsis* and tobacco. The Plant Cell 20, 1330–1345.

Illouz-Eliaz N, Nissan I, Nir I, Ramon U, Shohat H, Weiss D. 2020. Mutations in the tomato gibberellin receptors suppress xylem proliferation and reduce water loss under water-deficit conditions. Journal of Experimental Botany 71, 3603–3612.

Illouz-Eliaz N, Ramon U, Shohat H, Blum S, Livne S, Mendelson D, Weiss D. 2019. Multiple gibberellin receptors contribute to phenotypic stability under changing environments. The Plant Cell 31, 1506–1519.

Kalladan R, Lasky JR, Chang TZ, Sharma S, Juenger TE, Verslues PE. 2017. Natural variation identifies genes affecting drought-induced abscisic acid accumulation in *Arabidopsis thaliana*. Proceeding of the National Academy of Sciences of the USA 114, 11536–11541.

Kooyers NJ. 2015. The evolution of drought escape and avoidance in natural herbaceous populations. Plant Science 234, 155–162.

Köster J, Rahmann S. 2012. Snakemake- a scalable bioinformatics workflow engine. Bioinformatics 28, 2520–2522. 10.1093/bioinformatics/bts480

Lata C, Prasad M. 2011. Role of DREBs in regulation of abiotic stress responses in plants. Journal of Experimental Botany 62, 4731–4748.

Li J, Sima W, Ouyang B, et al. 2012. Tomato SlDREB gene restricts leaf expansion and internode elongation by downregulating key genes for gibberellin biosynthesis. Journal of Experimental Botany 63, 6407–6420.

Love MI, Huber W, Anders S. 2014. Moderated estimation of fold change and dispersion for RNA-seq data with DESeq2. Genome Biology 15, 1–21. 10.1186/s13059-014-0550-8

Martin M. 2011. Cutadapt removes adapter sequences from high-throughput sequencing reads. European Molecular Biology network Journal 17, 10–12. 10.14806/ej.17.1.200

Nakano T, Suzuki K, Fujimura T, Shinshi H. 2006. Genome-wide analysis of the ERF gene family in *Arabidopsis* and rice. Plant Physiology 140, 411–432.

Nir I, Shohat H, Panizel I, Olszewski N, Aharoni A, Weiss D. 2017. The tomato DELLA protein PROCERA acts in guard cells to promote stomatal closure. The Plant Cell 29, 3186–3197.

Noctor G, Mhamdi A, Foyer CH. 2014. The roles of reactive oxygen metabolism in drought: not so cut and dried. Plant Physiology 164, 1636–1648.

Ray PM, Green PB, Cleland R. 1972. Role of turgor in plant cell growth. Nature 239, 163–164.

Sarkar T, Thankappan R, Mishra GP, Nawade BD. 2019. Advances in the development and use of DREB for improved abiotic stress tolerance in transgenic crop plants. Physiology and Molecular Biology of Plants 25, 1323–1334.

Shani E, Hedden P, Sun Tp. 2024. Highlights in gibberellin research: A tale of the dwarf and the slender. Plant Physiology 195, 111–134.

Shohat H, Cheriker H, Kilambi HV, Illouz-Eliaz N, Blum S, Amsellem Z, Tarkowská D, Aharoni A, Eshed Y, Weiss D. 2021. Inhibition of gibberellin accumulation by water deficiency promotes fast and long-term ‘drought avoidance’ responses in tomato. New Phytologist 232, 1985–1998.

Skirycz A, Inzé D. 2010. More from less: plant growth under limited water. Current Opinion in Biotechnology 21, 197–203.

Sun S, Yu JP, Chen F, Zhao TJ, Fang XH, Li YQ, Sui SF. 2008. TINY, a dehydration-responsive element (DRE)-binding protein-like transcription factor connecting the DRE- and ethylene-responsive element-mediated signaling pathways in *Arabidopsis*. Journal of Biological Chemistry 283, 6261–6271.

Sun Y, Qiao Z, Muchero W, Chen JG. 2020. Lectin Receptor-Like Kinases: The sensor and mediator at the plant cell surface. Frontiers in Plant Science 11, 596301. 10.3389/fpls.2020.596301

Tardieu F, Simonneau T, Muller B. 2018. The physiological basis of drought tolerance in crop plants: a scenario-dependent probabilistic approach. Annual Reviews of Plant Biology 69, 733–759.

Vaid N, Macovei A, Tuteja N. 2013. Knights in action: Lectin Receptor-Like Kinases in plant development and stress responses. Molecular Plant 6, 1405–1418.

Wilson K, Long D, Swinburne J, Coupland G. 1996. A Dissociation insertion causes a semidominant mutation that increases expression of TINY, an *Arabidopsis* gene related to APETALA2. The Plant Cell 8, 659–671.

Xie Z, Nolan T, Jiang H, Tang B, Zhang M, Li Z, Yin Y. 2019. The AP2/ERF transcription factor TINY modulates brassinosteroid-regulated plant growth and drought responses in *Arabidopsis*. The Plant Cell 31, 1788–1806.

Zhang H, Zhu J, Gong Z, Zhu JK. 2022. Abiotic stress responses in plants. Nature Reviews Genetics 23, 104–119.

Zhang Y, Immink R, Liu CM, Emons AM, Ketelaar T. 2013. The *Arabidopsis* exocyst subunit SEC3A is essential for embryo development and accumulates in transient puncta at the plasma membrane. New Phytologist 199, 74–88.

